# Persistent immune imprinting after XBB.1.5 COVID vaccination in humans

**DOI:** 10.1101/2023.11.28.569129

**Authors:** M. Alejandra Tortorici, Amin Addetia, Albert J. Seo, Jack Brown, Kaitlin R. Sprouse, Jenni Logue, Erica Clark, Nicholas Franko, Helen Chu, David Veesler

## Abstract

Immune imprinting - also known as ‘original antigenic sin’ - describes how the first exposure to a virus shapes the immunological outcome of subsequent exposures to antigenically related strains. SARS-CoV-2 Omicron breakthrough infections and bivalent COVID-19 vaccination were shown to primarily recall cross-reactive memory B cells and antibodies induced by prior mRNA vaccination with the Wuhan-Hu-1 spike rather than priming naive B cells that recognize Omicron-specific epitopes. These findings underscored a strong immune imprinting resulting from repeated Wuhan-Hu-1 spike exposures. To understand if immune imprinting can be overcome, we investigated memory and plasma antibody responses after administration of the updated XBB.1.5 COVID mRNA vaccine booster. Our data show that the XBB.1.5 booster elicits neutralizing antibody responses against current variants that are dominated by recall of pre-existing memory B cells previously induced by the Wuhan-Hu-1 spike. These results indicate that immune imprinting persists even after multiple exposures to Omicron spikes through vaccination and infection, including post XBB.1.5 spike booster mRNA vaccination, which will need to be considered to guide the design of future vaccine boosters.

---

Emergence of immune evasive SARS-CoV-2 variants erodes the effectiveness of COVID-19 vaccines, which led to the development of two updated vaccine boosters. A bivalent Wuhan-Hu-1/BA.5 (or BA.1 for a few countries) spike (S) mRNA booster vaccine was approved in August 2022^1,2^ and, subsequently, a monovalent XBB.1.5 S mRNA booster vaccine in September 2023^3^. Neutralizing antibodies are a correlate of protection against COVID-19^4–7^ and most plasma neutralizing activity is directed to the S receptor-binding domain (RBD)^8–10^. Recent studies showed that antibody responses to Omicron variants are dominated by pre-existing immunity resulting from prior exposure to the Wuhan-Hu-1 spike, due to immune imprinting^11–15^. A study of individuals receiving inactivated Wuhan-Hu-1 viral vaccines, however, found that repeated Omicron infections could overcome immune imprinting, leading to elicitation of *de novo* antibody responses specific for these variants^16^. It is unknown if a similar outcome can be achieved through repeated administration of updated vaccine boosters in individuals previously imprinted via multiple Wuhan-Hu-1 S exposure.

To evaluate humoral immunity elicited upon receipt of an XBB.1.5 S mRNA vaccine booster, we collected plasma from individuals who had previously received multiple vaccine doses with or without known infection (**Tables S1-S2 in the Supplementary Appendix**). We used a vesicular stomatitis virus (VSV) pseudotyped with the Wuhan-Hu-1/D614G S, BQ.1.1 S, XBB.1.5 S or the BA.2.86 S to assess the potency and breadth of plasma neutralizing antibodies in this cohort and compare them with plasma collected upon receipt of the bivalent Wuhan-Hu-1/BA.5 vaccine booster. Neutralizing activity was highest against the Wuhan-Hu-1/G614 S VSV with geometric mean titers (GMTs) of 1,700 and 3,000 after receiving the XBB.1.5 S and the Wuhan-Hu-1/BA.5 S bivalent vaccine boosters, respectively. GMTs against BQ.1.1, XBB.1.5 and BA.2.86 S VSV were 510, 540 and 480 (after XBB.1.5 S vaccination) or 420, 240 and 320 (after bivalent vaccination), respectively. (**Figure 1A-B, Figure S1 in the Supplementary Appendix**). Our data show that the updated vaccine booster elicits neutralizing antibody responses against current variants, which is expected to translate into enhanced protection in the real world.

**Figure 1.**
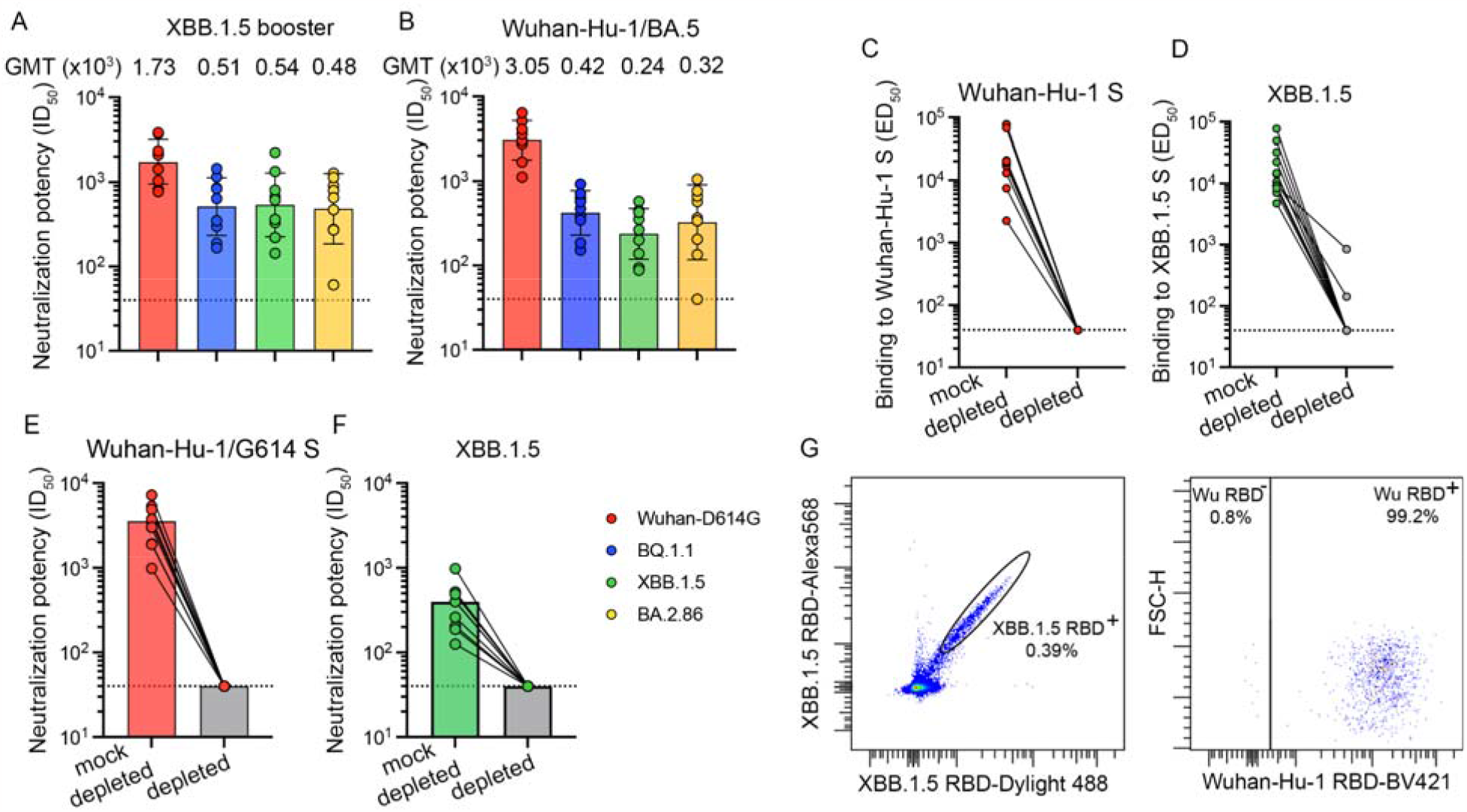
Immune imprinting dominates the immune response elicited upon XBB.1.5 S mRNA booster vaccination in humans. Panels A and B show plasma neutralizing antibody titers evaluated using a vesicular stomatitis virus (VSV) pseudotyped with the Wuhan-Hu-1 S harboring the D614G mutation, the BQ.1.1 mutations, the XBB.1.5 mutations or the BA.2.86 mutations using plasma obtained 7 to 13 days (mean: 9.7 days) after vaccination with the XBB.1.5 S mRNA booster (A) or 22 to 51 days (mean: 33 days) after vaccination with the bivalent Wuhan-Hu-1/BA.5 S mRNA booster. Geometric mean titers (GMTs) against each pseudovirus are indicated above the plot. Panels C and D show antibody binding titers (expressed as mean effective dilution 50%, ED_50_) against Wuhan-Hu-1 S (C) and XBB.1.5 S (D) before (left) and after (right) depletion of antibodies recognizing Wuhan-Hu-1 S, as determined by ELISA using XBB.1.5 S vaccinee plasma. Panels E and F show neutralizing antibody titers against and XBB.1.5 S (E) and Wuhan-Hu-1 S VSV (F) mock-depleted (left column) and after depletion (right column) of antibodies recognizing Wuhan-Hu-1 S. The dotted lines indicate the limit of detection of 1/40 for all assays. Data shown are from one representative (out of two) biological replicates with each point corresponding to the average of two technical replicates. Panel G shows the analysis of XBB.1.5 and Wuhan-Hu-1 RBD binding of memory B cells obtained from peripheral blood of individuals who received the XBB.1.5 S mRNA booster using flow cytometry.

The finding that administration of an XBB.1.5 S booster elicited higher plasma neutralizing activity against Wuhan-Hu-1/D614G S VSV (vaccine-mismatched) relative to XBB.1.5 S VSV (vaccine-matched) is a serological indication of immune imprinting. To investigate this further, we depleted polyclonal plasma antibodies recognizing the Wuhan-Hu-1 S trimer and assessed binding titers against the Wuhan-Hu-1 S and XBB.1.5 S ectodomain trimers using ELISAs. As expected, no antibodies binding to Wuhan-Hu-1 S were detected after depletion. Moreover, depletion of antibodies targeting Wuhan-Hu-1 S entirely abrogated binding to XBB.1.5 S except for two individuals for which binding titers decreased 10-fold (**Figure 1C-D, Figure S2 in the Supplementary Appendix**). Accordingly, we did not detect neutralizing antibodies against Wuhan-Hu-1/D614G S VSV and XBB.1.5 S VSV following depletion (**Figure 1E-F, Figure S3 in the Supplementary Appendix**), indicating the absence of XBB.1.5 S-specific antibodies in the plasma of these subjects (i.e. that were not cross-reactive with Wuhan-Hu-1 S). These data suggest that XBB.1.5 S vaccination increased cross-reactive plasma antibody titers previously elicited by Wuhan-Hu-1 S exposure, which are also binding to and neutralizing XBB.1.5 and other variants, instead of inducing *de novo* antibody responses against XBB.1.5 S.

We subsequently analyzed memory B cell populations found in the peripheral blood upon XBB.1.5 S vaccination by measuring the frequency of XBB.1.5 RBD-reactive memory B cells which also bound to the Wuhan-Hu-1 RBD using flow cytometry. Only 5 of the 12 individuals profiled had memory B cells that recognized the XBB.1.5 RBD, but not the Wuhan-Hu-1 RBD, and these memory B cells were rare (0.4-13.3%) (**Figure 1G, Figures S4-S5 and Table S1 in the Supplementary Appendix**). Given that the vast majority of XBB.1.5 RBD-binding memory B cells also bound to the Wuhan-Hu-1 RBD, *de novo* elicitation of memory B cells is possible through variant-specific mRNA booster, as is the case here for XBB.1.5, but difficult to induce due to preferential recall of pre-existing Wuhan-Hu-1 memory B cells.

The lack of detectable plasma antibodies specific for XBB.1.5 S and the scarcity of memory B cells binding to the XBB.1.5 RBD, but not the Wuhan-Hu-1 RBD, indicate that the humoral immune responses elicited by XBB.1.5 S vaccination are dominated by recall of pre-existing memory B cells previously induced by Wuhan-Hu-1 S vaccination instead of inducing *de novo* responses against this new variant. These findings concur with observations made after Omicron BA.1, BA.2 and BA.5 breakthrough infections^11,13^ and with that made after the roll out of the bivalent Wuhan-Hu-1/BA.5 and Wuhan-Hu-1/BA.1 S vaccine boosters^12,15^. As SARS-CoV-2 S-specific memory B cells continue to evolve and increase in frequency several months post infection or vaccination^17–21^, future analysis at later time points will shed light on long-term immunological impact of imprinting. Collectively, our results indicate that the updated XBB.1.5 S vaccine elicit neutralizing antibodies against circulating variants and point to the persistence of immune imprinting which will need to be considered to guide the design of future vaccine boosters.

## Acknowledgements

This study was supported by the National Institute of Allergy and Infectious Diseases (DP1AI158186, P01AI167966 and 75N93022C00036 to D.V.), a Pew Biomedical Scholars Award (D.V.), an Investigators in the Pathogenesis of Infectious Disease Awards from the Burroughs Wellcome Fund (D.V.), Fast Grants (D.V.), the University of Washington Arnold and Mabel Beckman Cryo-EM Center and the National Institute of Health grant S10OD032290 (to D.V.). D.V. is an Investigator of the Howard Hughes Medical Institute and the Hans Neurath Endowed Chair in Biochemistry at the University of Washington.

## Supplementary Methods

### Cell lines

Cell lines used in this study were obtained from JCRB-Cell Bank (VeroE6-TMPRSS2, geneticin resistant) and ThermoFisher Scientific (Expi293F™ cells). Cells were cultivated at 37°C, in an atmosphere of 5 % CO_2_ and with 130 rpm of agitation for suspension cells. None of the cell lines used were routinely tested for mycoplasma contamination. The VeroE6-TMPRSS2 puromycin resistant cells were previously described^22^.

### Plasmids

Plasmids encoding Wuhan-Hu-1 S Hexapro ectodomain (residues 1-1208) and XBB.1.5 S Hexapro ectodomain residues (residues 1-1203) were synthesized into pCDNA 3.1 (-) and pcDNA 3.1 (+), respectively. Both genes were synthesized by Genscript and harbor the HexaPro mutations^23^, a wildtype signal peptide, a furin cleavage site mutated _685_RSV_687_ to _685_SSV_687_, an avi-tag, and an octa-his tag for affinity purification.

Constructs for membrane-anchored S glycoproteins from SARS-CoV-2 Wuhan-D614G, BQ.1.1, XBB.1.5 and BA.2.86 contain the wild-type signal peptide and a 21-amino acid C-terminal deletion^24^ and were all synthesized and cloned into HDM vector. Genes were synthesized by Genscript with codon optimized for expression in mammalian cells, without tag and cloned in frame with a Kozak’s sequence to direct translation.

Plasmids encoding the SARS-CoV-2 Wuhan-Hu-1 and the XBB.1.5 RBDs were previously described elsewhere^12,25^. The SARS-CoV-2 Wuhan-Hu-1 RBD construct contains an N-terminal mu-phosphatase signal peptide and C-terminal octa-histidine tag followed by an avi-tag. The XBB.1.5 RBD construct contains an N-terminal BM40 signal peptide and a C-terminal octa-histidine tag followed by an avi-tag. The boundaries of both the Wu and XBB.1.5 RBD constructs are N-_328_RFPN_331_ and C-_528_KKST_532_.

### Recombinant protein production

SARS-CoV-2 Wuhan-Hu-1 and XBB.1.5 HexaPro S ectodomains were expressed in Expi293 cells at 37°C and 8% CO_2_. Cells were transfected with the corresponding plasmids using Expifectamine following the manufacturer’s instructions. Four days post-transfection, supernatants were clarified by centrifugation at 4121g for 30 minutes, supplemented with 25 mM phosphate pH 8.0, 300 mM NaCl, and 0.5 mM phenylmethylsulfonyl fluoride (PMSF). Supernatant was then 0.22µm vacuum filtered and passed through 1 mL His trap HP or Excel column (Cytiva) previously equilibrated in 25 mM phosphate pH 8.0, 300 mM NaCl. S proteins were eluted using a buffer identical to the binding buffer with the addition of 300 mM imidazole. Fractions containing the proteins were pooled and buffer exchanged to 50 mM Tris-HCl pH 8.0, 150 mM NaCl and stored at 4°C or immediately used.

The SARS-CoV-2 Wu and XBB.1.5 RBD proteins were expressed and purified as described above. Following buffer exchange, the purified RBDs were biotinylated using the BirA biotin-protein ligase reaction kit (Avidity). The biotinylated proteins were passed, washed, and eluted again on the same affinity column, concentrated and ran over a Superdex200 increase 10/300 size-exclusion column (Cytiva). Fractions corresponding to monomeric and monodisperse RBDs were collected, flash frozen, and stored at -80°C until use.

### Plasma antibody depletion

Invitrogen His-Tag Dynabeads (ThermoFisher 10104D) were used for depletion of plasma samples from antibodies recognizing the Wuhan-Hu-1 S trimer, as previously described^9^ with some modifications. Vortexed beads were incubated at room temperature on an Invitrogen DynaMag-2 Magnet (ThermoFisher 12–321-D) for two minutes to allow beads to separate for the liquid phase. Supernatant was discarded and beads were washed one time with TBS-T and divided in two tubes. After a two-minute incubation on the magnet, TBS-T supernatants were discarded and one set of beads was incubated with 4 mg of purified his-tagged SARS-CoV-2 Wuhan-Hu-1 S ectodomain trimer (Wuhan-Hu-1 S depletion) and the other set was incubated with TBS-T alone (mock depletion) with gentle rotation for 1 h at room temperature. Supernatants were discarded using the magnet and beads were washed three times with TBS-T. Subsequently, 20 µl of each of the plasma samples were incubated with 80 µl of S-loaded beads or mock-loaded beads for 1 h at 37°C during which time they were mixed every 15 min. Plasma samples were recovered using the magnet to separate the beads and used for neutralization assays.

### VSV pseudotyped virus production

Vesicular stomatitis virus (VSV) were pseudotyped with the SARS-CoV-2 S proteins corresponding to Wuhan-D614G, BQ.1.1, XBB.1.5 and BA.2.86 variants following a previously described protocol^26^. Briefly, HEK293T cells seeded in poly-D-lysine coated 10-cm dishes in DMEM supplemented with 10% FBS and 1% PenStrep were transfected with a mixture of 24 µg of the corresponding plasmids, 60 µl Lipofectamine 2000 (Life Technologies) in 5 ml of Opti-MEM, following the manufacturer’s instructions. After 5 h at 37°C, 5 ml of DMEM supplemented with 20% FBS and 2% PenStrep were added. The next day, cells were washed three times with DMEM and were transduced with VSVΔG-luc^27^. After 2 h, the virus inoculum was removed and cells were washed five times with DMEM prior to the addition of DMEM supplemented with anti-VSV-G antibody [Il-mouse hybridoma supernatant diluted 1 to 25 (v/v), from CRL-2700, ATCC] to minimize parental background. After 18-24 h, supernatants containing pseudotyped VSV were harvested, centrifuged at 2,000 x g for 5 min to remove cellular debris, filtered with a 0.45 µm membrane, concentrated 10 times using a 30 kDa cut off membrane (Amicon), aliquoted, and frozen at -80°C.

### Pseudotyped VSV neutralization

VeroE6-TMPRSS2 cells were seeded into coated clear bottom white walled 96-well plates at 40,000 cells/well and cultured overnight at 37°C. Eleven 3-fold serial dilutions of each plasma sample were prepared in DMEM. Pseudotyped VSV viruses, diluted 1 to 20 in DMEM containing anti-VSV-G antibody, were added 1:1 (v/v) to each plasma sample dilution and mixtures of 50 µl volume were incubated for 45-60 min at 37°C. VeroE6-TMPRSS2 cells were washed three times with DMEM and 40 μL of the mixture containing pseudotyped virus and plasma samples were added. Two hours later, 40 μL of DMEM were added to the cells. After 17-20 h, 70 μL of One-Glo-EX substrate (Promega) were added to each well and incubated on a plate shaker in the dark at 37°C. After 5-15 min incubation, plates were read on a Biotek Neo2 plate reader. Measurements were done in duplicate or triplicate with at least two biological replicates. Relative luciferase units were plotted and normalized in Prism (GraphPad): cells without pseudotyped virus added were defined as 0 % infection or 100 % neutralization, and cells with virus only (no plasma) were defined as 100 % infection or 0 % neutralization.

### Enzyme-linked immunosorbent assays (ELISA)

Analysis of plasma binding antibodies for samples mock-depleted or depleted of antibodies binding to the Wuhan-Hu-1 S ectodomain trimer was performed using ELISAs. Briefly, clear flat bottom Immuno Nonsterile 384-well plates (Thermo Scientific) were coated overnight at room temperature with 30 μl of Wuhan-Hu-1 S or XBB.1.5 S prepared at 3 μg/ml in PBS (137 mM of NaCl, 2.7 mM of KCl, 10 mM of Na2HPO4, and 1.8 mM of KH2PO4, pH 7.2). The next day, plates were blocked with Blocker™ Casein (Thermo Scientific) and subsequently incubated with serial dilutions of plasma samples for 1 h at 37°C. After four washing steps with TBS-T, goat anti-human IgG-Fc secondary antibody conjugated to HRP (Invitrogen A18817, diluted 1/500) was added and incubated for 1 h at 37°C. Plates were washed four times with TBS-T and KPL SureBlue Reserve™ TMB Microwell Peroxidase Substrate (VWR) was added. After 2 min incubation, 1N HCl was added and absorbance at 405 nm was measured using a Biotek Neo2 plate reader. Data were plotted using GraphPad Prism 9.1.0.

### Flow cytometry analysis of SARS-CoV-2 RBD-reactive memory B cells

To define specific B cell populations reactive with the XBB.1.5 and the Wuhan-Hu-1 RBDs, RBD–streptavidin tetramers conjugated to fluorophores were generated by incubating biotinylated RBDs with streptavidin at a 4:1 molar ratio for 30 min at 4 °C. Excess of free biotin was then added to the reaction to bind any unconjugated sites in the streptavidin tetramers. The RBD-streptavidin tetramers were washed once with PBS and concentrated with a 100-kDa cut-off centrifugal concentrator (Amicon). An additional streptavidin tetramer conjugated to biotin only was generated and included in the staining as decoy.

Approximately 5 to 15 million PMBCs were collected 7 to 13 days post-vaccination for individuals who received an XBB.1.5 S mRNA vaccine booster. Cells were collected by centrifugation at 400g for 2 mins at 4°C, washed twice with PBS and stained with Zombie Aqua dye (Biolegend; Catalog No. 423101; diluted 1:100 in PBS) at room temperature. After 30 min incubation, cells were washed twice with PBS and stained with antibodies for CD20-PECy7 (BD; Catalog No. 335793), CD3-Alexa eFluor780 (Thermo Fisher; Catalog No. 47-0037-41), CD8-Alexa eFluor780 (Thermo Fisher; Catalog No. 47-0086-42), CD14-Alexa eFluor780 (Thermo Fisher; Catalog No. 47-0149-42), CD16-Alexa eFluor780 (Thermo Fisher; Catalog No. 47-0168-41), IgM-Alexa Fluor 647 (BioLegend; Catalog No. 314535), IgD-Alexa Fluor 647 (BioLegend; Catalog No. 348227), and CD38-Brilliant Violet 785 (BioLegend; Catalog No. 303529), all diluted 1:200 in Brilliant Stain Buffer (BD; Catalog No. 563794), along with the RBD-streptavidin tetramers for 30 min at 4°C. Cells were washed three times, resuspended in PBS, and passed through a 35-μm filter before being examined on a BD FACSymphony A3 for acquisition and FlowJo 10.8.1 for analysis. Gates for identifying the XBB.1.5 RBD double-positive population as well as the subsequent Wuhan-Hu-1 RBD-positive and Wuhan-Hu-1 RBD-negative populations were drawn based on staining of fluorescent minus one controls.

---

**Figure S1.**
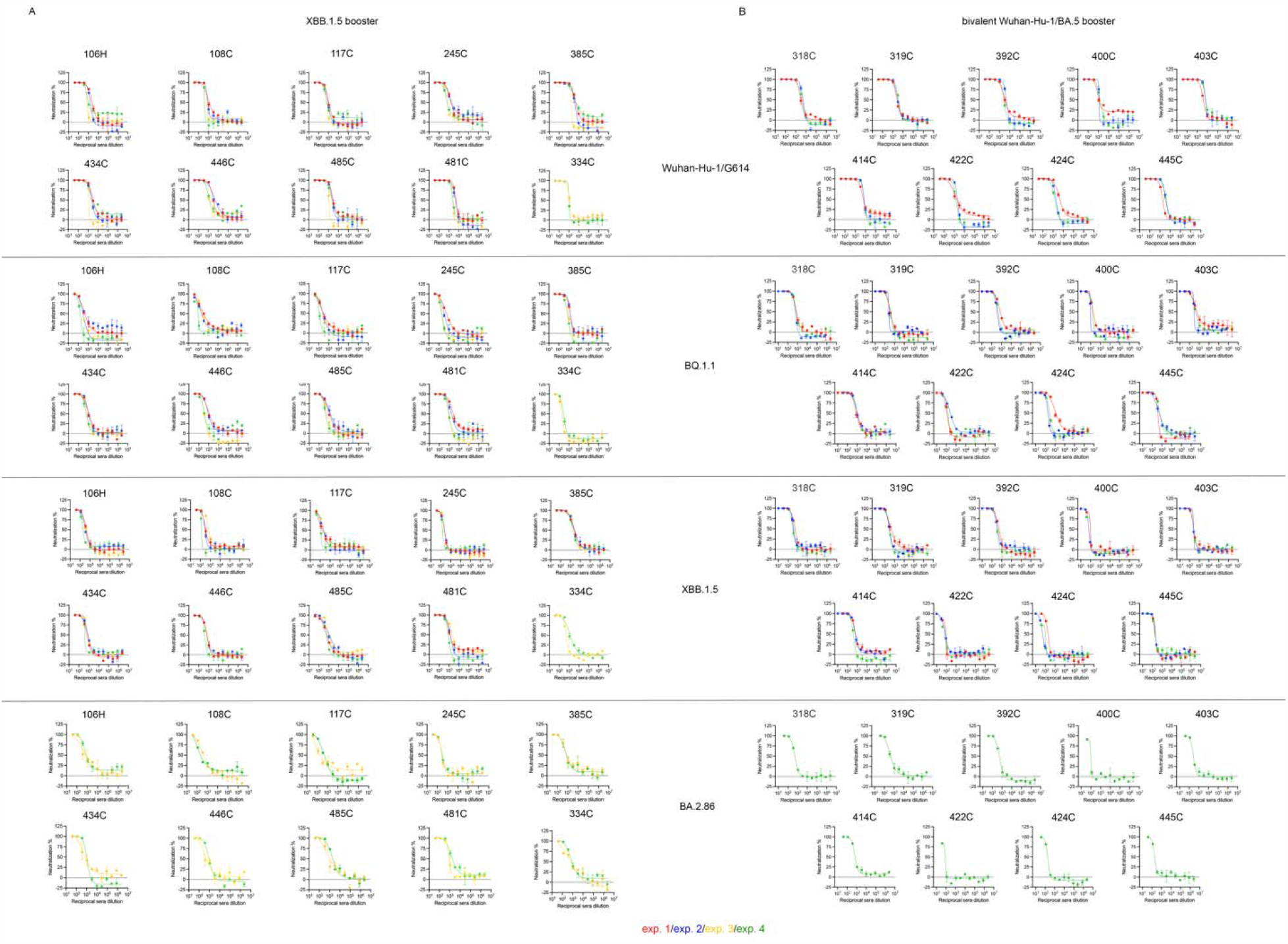
Plasma neutralizing antibody titers after vaccination with the XBB.1.5 S mRNA booster (A) and after the bivalent Wuhan-Hu-1/BA.5 S mRNA booster (B). Dose-response curves for all repeats are shown with each patient ID indicated on top of the graphs. Experiments 1 and 2 (exp. 1 and exp. 2) were performed using a first batch of VSV S pseudotyped viruses using VeroE6-TMPRSS2 cells puromycin resistant. Experiments 3 and 4 (exp. 3 and exp 4) were performed using a second batch of VSV S pseudotyped viruses using VeroE6-TMPRSS2 cells geneticin resistant for exp. 3 and VERO-TMPRSS2 cell puromycin resistant for exp. 4.

**Figure S2.**
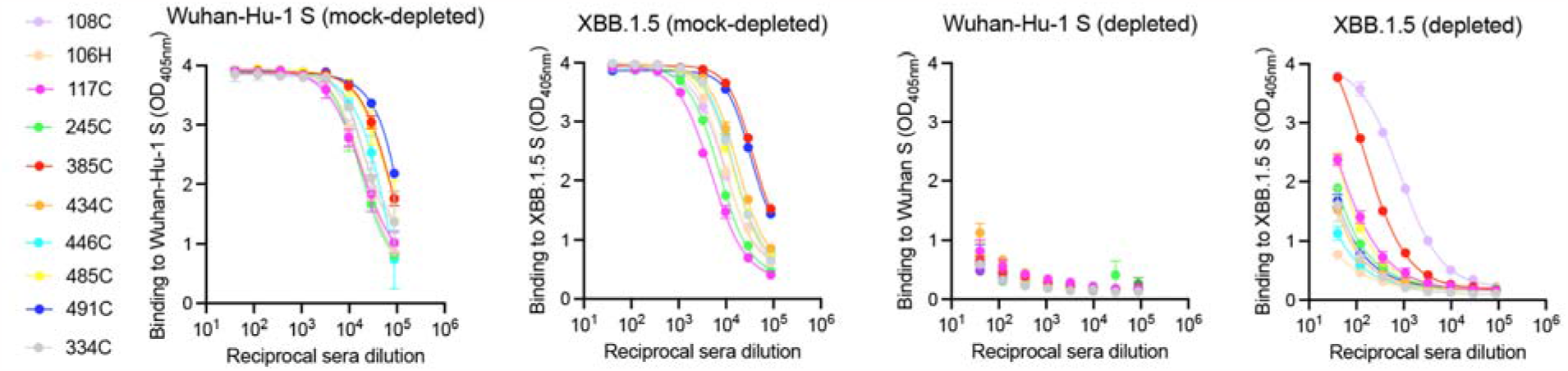
Plasma antibody binding titers against Wuhan-Hu-1 S and XBB.1.5 S after vaccination with the XBB.1.5 S mRNA booster in plasma samples mock-depleted and depleted of Wuhan-Hu-1 S-directed antibodies. Dose-response curves for one representative experiment are shown. The color key indicates patient IDs.

**Figure S3.**
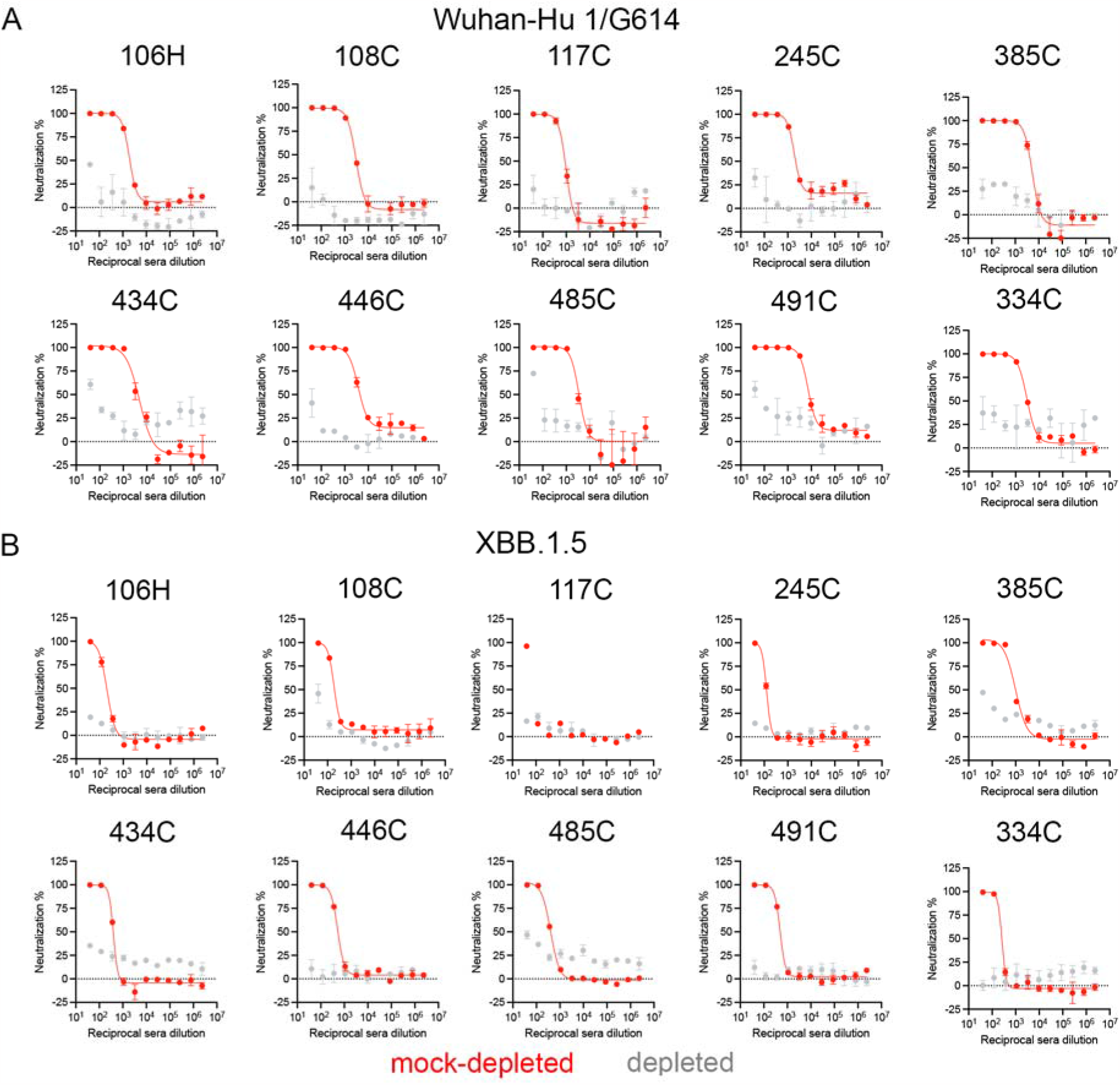
Plasma neutralizing antibody titers after vaccination with the XBB.1.5 S mRNA booster in plasma samples mock-depleted and depleted of Wuhan-Hu-1 S-directed antibodies. Dose-response curves for one representative experiment are shown. Patient IDs are indicated on top of each graph.

**Figure S4.**
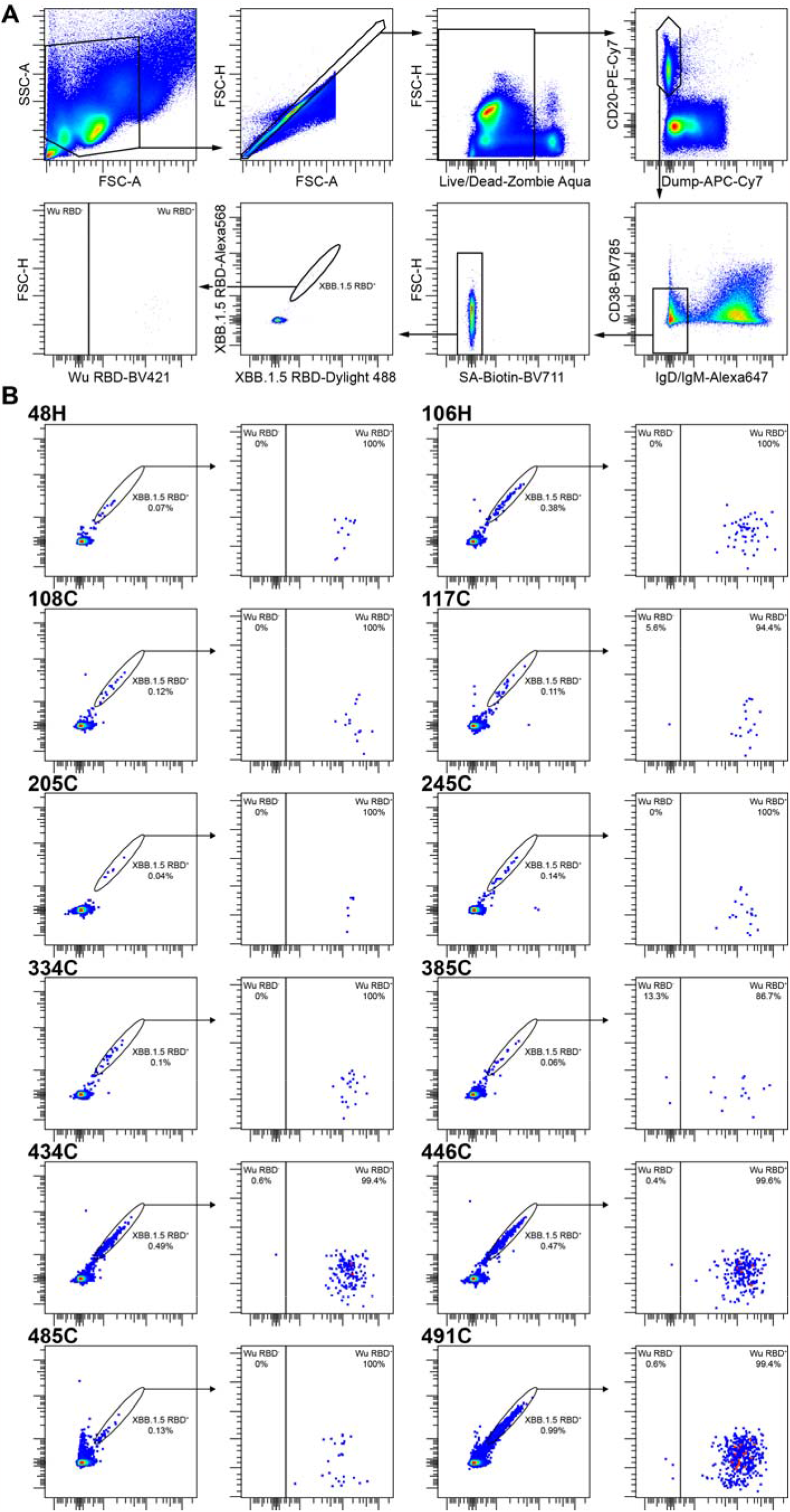
Flow cytometry analysis of memory B cells. Panel A shows the gating strategy to evaluate the cross-reactivity with the Wuhan-Hu-1 RBD of XBB.1.5 RBD+ memory B cells. Dump includes markers for CD3, CD8, CD14, and CD16. Panel B shows the gating of XBB.1.5 RBD-reactive memory B cells and subsequent Wuhan-Hu-1 RBD-binding of these memory B cells for each individual profiled.

**Table S1.** Demographics information of the subjects in the XBB.1.5 vaccinee cohorts.

| Patient ID | Number of pre-Omicron SARS-CoV-2 infection (before 11/2021) | Number of post-Omicron SARS-CoV-2 infection (after 11/2021) | Number of Wuhan-Hu-1 vaccine doses | Number of bivalent Wuhan-Hu-1/BA.5 vaccine doses | Number of XBB.1.5 vaccine doses | Days between XBB.1.5 vaccine and blood draw |
| --- | --- | --- | --- | --- | --- | --- |
| 106H | 0 | 0 | 3 | 1 | 1 | 7 |
| 108C | 1 | 1 | 4 | 2 | 1 | 8 |
| 117C | 1 | 0 | 4 | 1 | 1 | 10 |
| 245C | 1 | 1 | 3 | 1 | 1 | 9 |
| 334C | 0 | 1 | 3 | 1 | 1 | 13 |
| 385C | 1 | 0 | 3 | 1 | 1 | 11 |
| 434C | 0 | 1 | 4 | 1 | 1 | 11 |
| 446C | 0 | 1 | 4 | 0 | 1 | 13 |
| 485C | 0 | 1 | 4 | 1 | 1 | 11 |
| 491C | 0 | 1 | 4 | 1 | 1 | 10 |

**Table S2.** Demographics information of the subjects in the Wuhan-Hu-1/BA.5 bivalent vaccinee cohorts.

| Patient ID | Number of pre-Omicron SARS-CoV-2 infection (before 11/2021) | Number of post-Omicron SARS-CoV-2 infection (after 11/2021) | Number of Wuhan-Hu-1 vaccine doses | Number of bivalent Wuhan-Hu-1/BA.5 vaccine doses | Number of XBB.1.5 vaccine doses | Days between bivalent Wuhan-Hu-1/BA.5 vaccine and blood draw |
| --- | --- | --- | --- | --- | --- | --- |
| 318C | 0 | 1 | 3 | 1 | 0 | 43 |
| 319C | 0 | 1 | 3 | 1 | 0 | 29 |
| 392C | 0 | 1 | 5 | 1 | 0 | 41 |
| 400C | 0 | 1 | 3 | 1 | 0 | 23 |
| 403C | 0 | 1 | 3 | 1 | 0 | 34 |
| 414C | 0 | 1 | 3 | 1 | 0 | 52 |
| 422C | 0 | 1 | 3 | 1 | 0 | 37 |
| 424C | 0 | 1 | 3 | 1 | 0 | 30 |
| 445C | 0 | 1 | 3 | 1 | 0 | 37 |

